# A canonical chloroplast unfolded protein response triggered by misfolded polypeptides

**DOI:** 10.1101/2025.06.11.659047

**Authors:** Alejo Cantoia, Fabricio Bertero, Rodrigo Berrocal, Eduardo A. Ceccarelli, Nicolás E. Blanco, Germán L. Rosano

## Abstract

Photosynthetic efficiency and plant viability rely on chloroplast protein homeostasis. While unfolded protein responses (UPRs) in the endoplasmic reticulum and mitochondria have been extensively characterized, the chloroplast UPR (cpUPR) remains less defined, partly due to the off-target effects of traditional stress-inducing methods. In this study, we provide direct evidence for the existence of the cpUPR by expressing engineered, folding-defective variants of ferredoxin-NADP⁺ reductase (FNR) in plant chloroplasts. The expression of aggregation-prone proteins inside the organelle triggered a robust upregulation of chloroplast quality control components, including CLPB3, CLPC1/C2, and HSP90C, as revealed by immunodetection and quantitative proteomics. The proteomic response scaled with the severity of the folding defect, with the fully insoluble FNR Δ20 variant inducing broader changes than the partially soluble Δ3 variant. Network analysis showed that most differentially abundant proteins were chloroplastic and clustered into functional groups related to proteostasis and photosynthesis. Comparative analysis with lincomycin-treated plants highlighted the specificity and advantages of using misfolded proteins to study the cpUPR. Expression of the folding variants conferred tolerance to heat treatment, suggesting that the response can enhance plant fitness. Taken together, our work establishes the cpUPR as a specific stress response in chloroplasts and provides new tools for its characterization.

## INTRODUCTION

Protein homeostasis, or proteostasis, refers to the balance between protein synthesis, folding, and degradation. Reaching a proper three-dimensional conformation is essential for protein solubility and function, and misfolding and aggregation can severely disrupt cellular physiology. Proteostasis is challenged by intrinsic factors such as errors in gene expression and by external stresses. Among these, abiotic stresses like elevated or freezing temperatures are particularly damaging, as they can destabilize protein structures, leading to misfolding, loss of function, and aggregation (Mogk et al., 2018). Cells can cope with the unfolded protein challenge by activating molecular responses that counterbalance the pathologic state. These pathways constitute the “unfolded protein response” (UPR), which comprises the activation of sensors and stress regulators that orchestrate an increase in the synthesis of proteins involved in quality control mechanisms, such as molecular chaperones and proteases (Hetz, 2012; Read & Schröder, 2021). The UPR is typically compartmentalized, and its molecular intricacies depend on the specific organelle affected. It was first discovered in the endoplasmic reticulum (Kozutsumi et al., 1988), where it is activated by the accumulation of misfolded proteins in the lumen under stress (erUPR). Similar responses were later found in the cytosol (Metzger & Michaelis, 2009; Wu, Meyer, Richter, et al., 2019) and other organelles, such as mitochondria (mtUPR) (Hoogenraad, 2017).

By its very definition, the UPR is triggered by the buildup of unfolded proteins within a cellular compartment. This state can be achieved in several ways. For instance, chemicals that interfere directly or indirectly with protein folding lead to the accumulation of unfolded or misfolded proteins. As an example, tunicamycin inhibits the initial stage of N-linked glycan production in the endoplasmic reticulum of eukaryotic cells, resulting in numerous misfolded proteins (Heifetz et al., 1979). In contrast, reducing agents like dithiothreitol impair oxidative protein folding (Braakman et al., 1992; Gopakumar & Singh, 2022). Cell treatment with these agents unleashes the erUPR (reviewed in Chapman et al., 1998). In mitochondria, treatments with paraquat (an inhibitor of the mitochondrial complex I that increases levels of reactive oxygen species) or CDDO (an inhibitor of the mitochondrial Lon1 protease) induce a potent mitochondrial unfolded protein response (mtUPR) (Münch & Harper, 2016; Shpilka & Haynes, 2018; Yoneda et al., 2004). These chemicals target critical enzymes involved in protein folding and degradation. For this reason, knockout organisms in the respective genes have also been extensively used to cause the accumulation of misfolded proteins (Shpilka & Haynes, 2018). On a larger scale, organismal-level abiotic stresses (e.g., heat shock) can also disrupt protein structure, generating responses with similar characteristics of a prototypical UPR.

Nevertheless, cell treatment with protein-folding disruptive chemicals or the use of knockout (or knockdown) organisms can activate other overlapping mechanisms, thereby obscuring the actual molecular signature of the UPR. A significant body of work suggests that pleiotropic effects in the chemically induced erUPR are numerous and activate additional counteractions beyond the prototypical UPR (Bergmann et al., 2018; Bergmann & Molinari, 2018; Gopakumar & Singh, 2022). Considering these issues, the sole accumulation of misfolded proteins appears as an unbiased strategy to trigger a true UPR. This approach has been used to differentiate the mtUPR and the erUPR from other mechanisms. In the former case, Zhao et al. used a folding-defective version of the mitochondrial protein ornithine transcarbamylase lacking residues 30 to 114 (Q. Zhao et al., 2002). Its expression in mitochondria resulted in the upregulation of nuclear genes encoding chaperonin 60, chaperonin 10, mtDNAJ, and the protease CLPP, among others, leading to a significant increase in the levels of these proteins. The abundances of non-mitochondrial chaperones belonging to the erUPR (BiP, Grp94, protein disulfide isomerase, calnexin, and calreticulin) were not affected, indicating that accumulation of the folding-defective protein in mitochondria caused selective stress in the organelle (Q. Zhao et al., 2002). Using a similar approach, Hidvegi et al. employed HeLa cells that accumulated nonpolymerogenic versions of the ER-resident protein α1-antitrypsin, which lacked 19 residues from their C-terminus. An increase in the synthesis of the ER chaperone BiP was readily detected, a hallmark in the activation of the erUPR (Hidvegi et al., 2005).

In chloroplasts, a UPR (cpUPR) has been inferred from the use of chemicals, gene knockdowns, or systemic stresses that interfere with protein folding or degradation within the organelle. The cpUPR was first described in *Chlamydomonas reindhartii*, where an inducible system was devised to achieve reduced levels of the essential chloroplastic protease CLPP (Ramundo & Rochaix, 2014b). The damage is so extensive that this system ultimately leads to organelle autophagy. Later, it was found that lincomycin (LIN) treatment of *Arabidopsis thaliana* plantlets triggered a response with similar characteristics of a UPR (Llamas et al., 2017). LIN is an antibiotic that inhibits chloroplast translation by binding to plastid ribosomes. As ribosomes halt, nascent polypeptides cannot properly fold and aggregate. However, LIN also negatively affects the transcription rate of photosynthesis-related genes and decreases the phosphorylation level of the light-harvesting complex II (Mulo et al., 2003). Moreover, it can affect chloroplast biogenesis and development, as well as the expression of light- and hormone-responsive genes and the circadian rhythm (Ruckle et al., 2012). Finally, systemic abiotic stresses also trigger cpUPR-like mechanisms. High-light treatment of leaves causes photooxidative damage to the photosystem II reaction center proteins, such as D1 and D2. The impaired proteins are targeted and degraded by the thylakoid-anchored metalloprotease FTSH2, also called VAR2 (Kato et al., 2012). Inactivation of FTSH2 in *var2* mutants grown under continuous light leads to the accumulation of damaged proteins in chloroplasts. At the same time, it also results in the buildup of reactive oxygen species, which could then disturb the redox balance within the chloroplasts, magnifying the disruption of proteostasis. Both phenomena trigger a cpUPR-like response, termed cpUPR-like damaged protein response (DPR) (Dogra et al., 2019). Altogether, these pioneering works used chemicals, knockdown organisms, and more general stresses to unleash protective molecular responses in plastids. Alas, their off-target effects on other cellular compartments or molecular components activate overlapping or unspecific pathways, which cannot be distinguished from the actual cpUPR.

Our work introduces the novel application of folding-defective proteins as triggers for the chloroplast unfolded protein response. We used two variants of ferredoxin NADP^+^ reductase (FNR) engineered to display folding defects. FNR is a monomeric, highly soluble enzyme that contains noncovalently bound FAD as a prosthetic group. It catalyzes the electron transfer from ferredoxin, reduced by photosystem I, to NADP+. Its carboxyl-terminal region includes most of the NADP(H) binding residues, some of which also participate in the non-covalent binding of the FAD ligand. To evaluate the impact of the accumulation of folding variants in chloroplasts, we used two approaches: transient transformation of *Nicotiana benthamiana* leaves and stable transformation of *A. thaliana*. Through a multi-methodological approach spearheaded by a comprehensive proteomics analysis, we sought to unravel the network of proteins involved in the canonical cpUPR.

## MATERIALS AND METHODS

### Plasmid construction

The construction of *Pisum sativum* FNR variants with reduced solubility was described in Orellano et al. The FNR Δ3 variant lacks Asp289, Trp290, and Ile291 residues. In contrast, the Δ20 deletion variant (FNR Δ20) lacks the last 20 residues from the C-terminal end. Coding sequences of wild-type FNR (FNR WT) and the aggregation-prone forms were cloned into the plant expression vector pCHF3. The coding sequence of the transit peptide from *A. thaliana* FNR leaf isoform 1 from chloroplasts was added to each construct to direct the proteins into the organelle. For confocal microscopy imaging, the cyan fluorescence protein (CFP) coding sequence, which was codon optimized for plant expression, was inserted between the transit peptide and the FNR variants. All cloning procedures were performed using RF cloning, and plasmid construction was verified through DNA sequencing.

### Plant transformation

The pCHF3 plasmids were introduced into the *Agrobacterium tumefaciens* strain GV3101-pMP90 by electroporation. For transient expression assays, the transformed *A. tumefaciens* cells harboring each vector were cultured until the late exponential phase, then centrifuged and resuspended in infiltration buffer (10 mM MES, pH 5.0, 10 mM MgCl2, and 200 µM acetosyringone) to achieve an optical density of 0.7 at 600 nm. Subsequently, each culture was utilized for leaf infiltration in 4-week-old *N. benthamiana* plants (Blanco et al., 2019).

Following agroinfiltration, the plants were maintained under control conditions in a growth chamber with a 16-hour photoperiod at 23°C during the day and 18°C at night for varying post-infiltration (PI) durations as required by the experiment. To generate plants with stable expression of the FNR variants, inflorescences of *A. thaliana* (Col-0 ecotype) were subjected to agrotransformation using the floral dip method (Clough & Bent, 1998). Seeds obtained from the transformed plants were subjected to surface sterilization and subsequently placed on plates containing 0.5× Murashige-Skoog salts, 0.8% agar, and 50 µg/mL kanamycin for selection purposes.

### Chloroplast isolation

4-week-old *N. benthamiana* plants were agroinfiltrated, as previously explained. 72 hours PI, they were transferred to darkness for 24 hours. To begin, the leaves were washed with cold distilled water and then homogenized using a Polytron blender for 15 s in the presence of cold CIB buffer (50 mM HEPES, pH 7.5, 330 mM sorbitol, 2 mM EDTA, 1 mM MgCl2, 0.25% w/v bovine serum albumin, 100 mM glutathione). The homogenized material was filtered through two layers of Miracloth, and the liquid was collected in a beaker. This homogenization and filtration step was repeated five times. The suspension was centrifuged for 15 min at 4000 rpm in a refrigerated centrifuge. After removing the supernatant, the chloroplasts were resuspended in 2 mL of CIB buffer using a paintbrush. The suspension was centrifuged on a Percoll (Percoll Plus, GE) discontinuous gradient (40%/80% layers in CIB buffer) at 7500 rpm for 30 min using a swinging bucket rotor to enrich the solution with intact chloroplasts, which migrate at the 40%/80% Percoll interphase. The chloroplasts were then transferred to a clean tube for washing with HF buffer (50 mM HEPES, pH 7.5, 330 mM sorbitol). Finally, a liquid nitrogen shock was applied to rupture the chloroplasts, and the samples were resuspended in denaturing sample buffer (1X) for SDS-PAGE electrophoresis.

### High-temperature treatment of plants

28-day-old *A. thaliana* plants were subjected to heat stress treatment in controlled environmental chambers (Fitotron, 60% relative humidity). T0 generation transgenic lines expressing pea FNR WT or folding variants and a control line transformed with the empty vector (pCHF3) were grown under identical conditions and treated in parallel. The heat stress protocol consisted of a preconditioning step, during which plants were incubated for 12 h at 19°C in darkness, followed by a heat shock phase at 45°C for 12 h under constant illumination (100 µmol m⁻² s⁻¹). In parallel, a non-stressed control group was incubated for the same 12-hour period under identical light and humidity conditions but at 22°C to assess normal growth.

### Confocal microscopy

Laser confocal scanning microscopy was performed using a Plan Apochromat 20x, 0.8-NA lens on an LSM880 Zeiss microscope, with the 458-nm laser line for excitation. The fluorescence emission was collected between 463 and 517 nm for CFP and 666 and 735 nm for chlorophyll. Cellular parameters and fluorescence intensities were analyzed with Fiji (Schindelin et al., 2012).

### Western blot analysis

Homogenates were prepared from leaf tissue by pulverization in liquid nitrogen, followed by resuspension in lysis solution (100 mM Tris-HCl pH 8, 10 mM MgCl2, 25 mM DTT, 1 mM EDTA, 20% v/v glycerol, and 1 mM phenylmethylsulfonyl fluoride). The extract was centrifuged at 12,000 g and 4°C for 15 minutes, and the resulting supernatant was used for protein separation by electrophoresis and immunodetection assays. Total protein content was quantified using the Bradford method. The protein extracts were separated by SDS-PAGE in 10% polyacrylamide gels and then transferred to nitrocellulose sheets. During the Western blot procedure, primary antibodies were used at a dilution of 1:3000, including anti-FNR (rabbit, AS15 2909, Agrisera), anti-CLPC (rabbit, AS01 001, Agrisera), and anti-CLPB3 (Parcerisa et al., 2020). Different types of secondary anti-rabbit IgG antibodies were used. In assays using enzyme-conjugated antibodies, alkaline phosphatase-conjugated antibodies were used at a dilution of 1/8000. In other immunodetection assays, chemiluminescence detection was employed using HRP-conjugated antibodies from Bio-Rad and the commercial detection kit provided by the same supplier (Inmun-Star HRP).

### Sample preparation and LC-MS/MS analysis

Protein extracts prepared as described were subjected to SDS-PAGE, with electrophoresis carried out until the proteins had migrated approximately 1 cm into the resolving gel. The gel section was excised, and proteins were reduced with 10 mM dithiothreitol in 50 mM ammonium bicarbonate at 56 °C for 45 min, followed by alkylation with 55 mM iodoacetamide in 50 mM ammonium bicarbonate in the dark at room temperature for 30 min. Gel pieces were then washed, dehydrated with acetonitrile, and rehydrated with sequencing-grade trypsin (Promega) at a concentration of 12.5 ng/μL in 50 mM ammonium bicarbonate. Digestion was carried out overnight at 37 °C. Peptides were extracted with 100% v/v acetonitrile, followed by 0.5% v/v trifluoroacetic acid, dried by vacuum centrifugation, and resuspended in 1% v/v formic acid.

Peptide separations were conducted using a nano HPLC Ultimate3000 system (Thermo Scientific) equipped with a nano column (EASY-Spray ES903, 50 cm × 50 μm ID, PepMap RSLC C18). The mobile phase consisted of 0.1% formic acid in water (solvent A) and 0.1% formic acid in acetonitrile (solvent B) at a 300 nL/min flow rate. The gradient profile was programmed as follows: (i) for LIN treatment: 4 – 30% solvent B over 114 min, 30 – 80% solvent B over 14 min, and 80% solvent B for 2 min; (ii) for proteomic comparison of plants expressing FNR variants: 4 – 35% solvent B over 90 min, 35 – 90% solvent B over 20 min, and 90% solvent B for 5 min. Mass spectrometry analysis was performed using a Q-Exactive HF mass spectrometer (Thermo Scientific). Ionization was achieved with a liquid junction voltage of 1.9 kV and a capillary temperature of 250°C. The full scan method employed a mass range of m/z 375–1,600, with an Orbitrap resolution of 120,000 (at m/z 200). The automatic gain control (AGC) target was set to 2 × 10⁵, and the maximum injection time was 32 ms. After the survey scan, the 20 most intense precursor ions were selected for MS/MS fragmentation. Fragmentation was performed with a normalized collision energy of 27 eV, and MS/MS scans were acquired with a dynamic first mass. The AGC target for MS/MS was also 2 × 10⁵, with a resolution of 15,000 (at m/z 200). An intensity threshold of 2.5 × 10⁵ was applied, and an isolation window of 1.4 m/z units was used. The maximum injection time for MS/MS scans was 32 ms. To improve data quality, charge state screening was enabled to reject unassigned, singly charged ions and ions with eight or more charges. A dynamic exclusion time of 30 s was implemented to prevent the re-selection of previously analyzed ions.

### MS data analysis

MS data were analyzed using standardized workflows with Proteome Discoverer (V: 2.4.1.15). Mass spectra *.raw files were searched against the reviewed protein database from *A. thaliana* (Mouse-ear cress, cv Columbia), entry UP000006548 from UniProt. Precursor and fragment mass tolerances were set to 10 ppm and 0.02 Da, respectively, allowing for up to two missed cleavages. Fixed modification: carbamidomethylation of cysteines. Dynamic modifications: protein N-terminal acetylation and/or methionine loss and methionine oxidation. The mass spectrometry proteomics data have been deposited to the ProteomeXchange Consortium *via* the PRIDE partner repository with the dataset identifier PXD064779 (Perez-Riverol et al., 2025).

The normalized abundances were uploaded to the Perseus platform for quality control and statistical analysis (Tyanova et al., 2016). Network analysis was conducted in Cytoscape, utilizing the StringApp to retrieve protein-protein interaction information and Gene Ontology (GO) terms. Clustering was performed using the clusterMaker app, and the MCL algorithm was applied with default settings.

## RESULTS

### Expression of FNR Δ3 in chloroplasts increases the levels of the CLPB disaggregase

Our working hypothesis is that the accumulation of unfolded proteins in the chloroplast is sufficient to unleash the cpUPR. We chose ferredoxin-NADP+ reductase (FNR) from pea as a model. We have previously shown that deletions at the C-terminus are detrimental to catalysis and solubility. Eliminating the triad Asp289, Trp290, and Ile291 leads to a complete loss of catalytic activity and a 50% reduction in solubility compared to the wild-type enzyme (Orellano et al., 1993). The deletion mutant FNR Δ3 does not bind FAD, as evidenced by the lack of the characteristic yellowish color of recombinant FNR preparations from bacterial cells (Figure 1). A more extensive truncation, FNR Δ20, removes the final 20 amino acids, further reducing solubility and resulting in an entirely insoluble protein. This Δ20 variant will be used in subsequent experiments to assess the effects of expressing an unfolded protein in *A. thaliana*.

**Figure 1:**
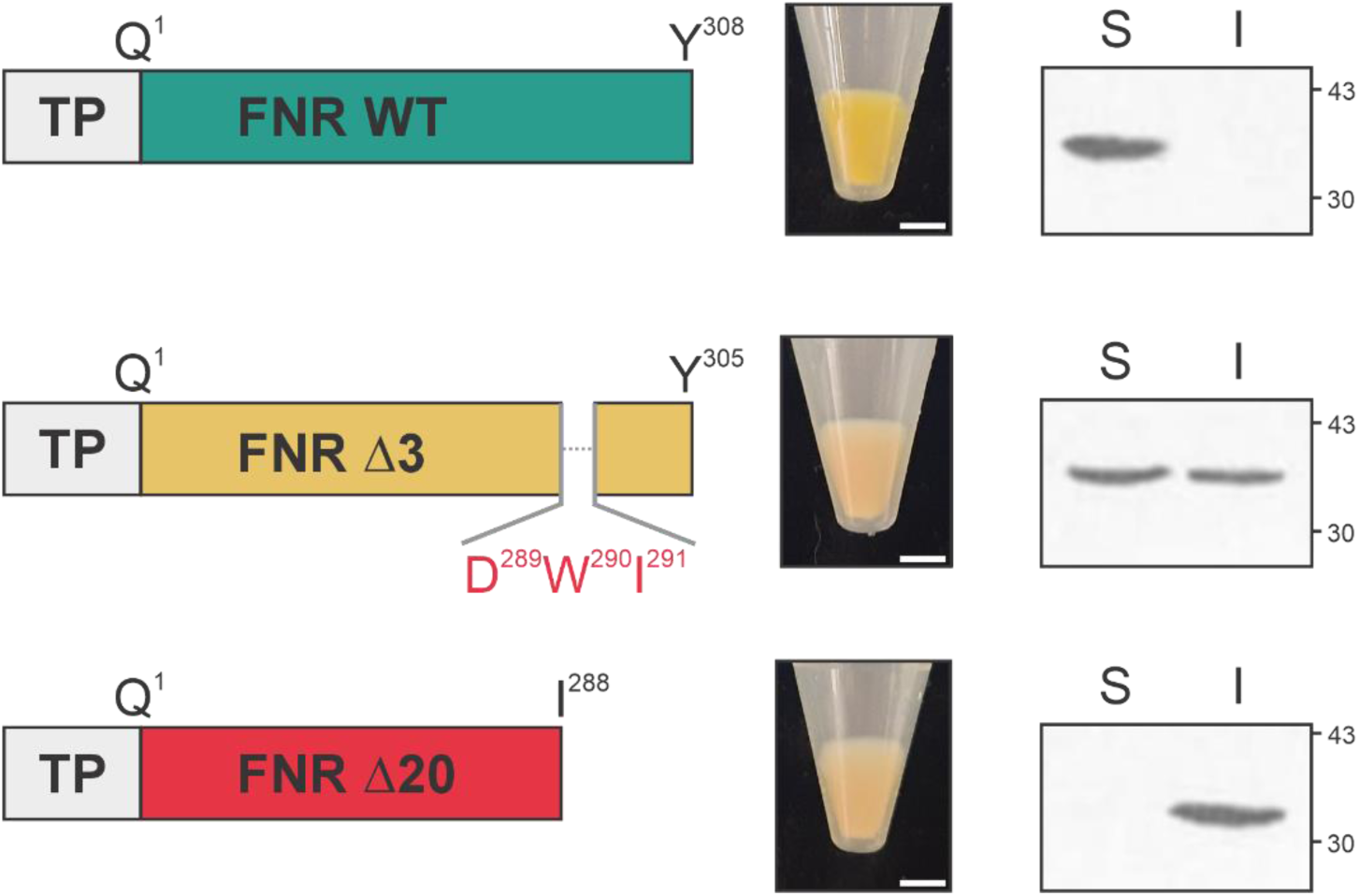
Expression and solubility of FNR C-terminal variants. Left: schematic representations of coding sequences for TP-FNR wild-type (WT), TP-FNR Δ3 (lacking residues D289–I291), and TP-FNR Δ20 (truncated at residue 288). Molecular mass of TP: 5.1 kDa. Middle: bacterial pellets show loss of FAD-dependent yellow color in both deletion variants. Scale bar: 0.5 cm. Right: Western blot of soluble (S) and insoluble (I) fractions. Numbers next to the blots represent molecular weight standards (in kDa).

The localization of the proteins to the chloroplast was achieved by fusing the transit peptide (TP) of *A. thaliana* FNR leaf isoform 1 from chloroplasts to the different pea FNR versions. This TP successfully directs proteins to the organelle in many plant species. Moreover, *in silico* analysis of the constructs by TargetP-2.0 indicated a high probability of chloroplast localization for all FNRs (Almagro Armenteros et al., 2019). TP-FNR WT and TP-FNR Δ3 coding sequences were cloned in the binary vector pCHF3 downstream of the 35S promoter for constitutive expression and used to transiently transform *N. benthamiana* leaves by agroinfiltration. Overexpression of FNR WT was confirmed by Western blot analysis of total *N. benthamiana* leaf extracts, showing detectable accumulation 48 hours post-infiltration, which persisted for at least four days (Figure 2A). The anti-FNR antibodies cross-react with the endogenous *N. benthamiana* FNR and the heterologous pea FNR WT (molecular mass of the mature forms: 34.6 and 34.8 kDa, respectively), as indicated by the signal observed in plant extracts transformed with the empty vector. The strong signal observed in plants overexpressing FNR WT reflects the combined contribution of endogenous and exogenous FNR. However, FNR Δ3 does not noticeably increase the intensity of the comigrating band, suggesting it is much less expressed than FNR WT despite differing by only three amino acids. When detection was performed on chloroplast-enriched fractions, faint expression of FNR Δ3 could be visualized as a band slightly more intense than in plants infiltrated with the empty vector (Figure 2A). Notably, no precursor forms of either protein were detected, suggesting that both FNR WT and FNR Δ3 were efficiently imported into chloroplasts and processed to their mature forms.

**Figure 2.**
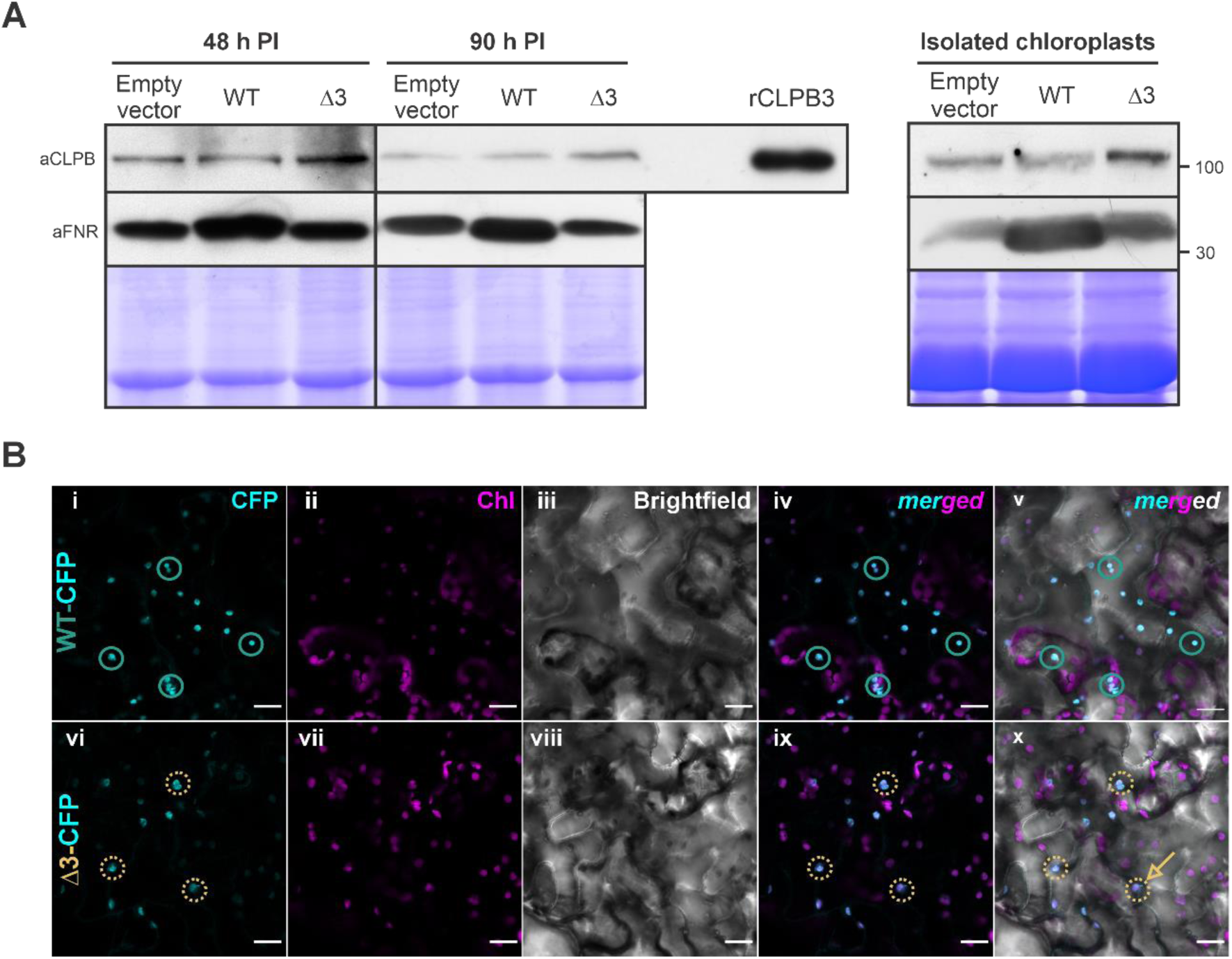
Expression and localization of pea FNRs and CLPB induction by FNR Δ3 in *N. benthamiana*. (A) Total protein extracts and chloroplast-enriched fractions from *N. benthamiana* leaves transiently expressing plastid-targeted FNR WT or FNR Δ3 were analyzed by immunoblotting at 48- and 90-hours post-infiltration (h PI). Antibodies against FNR (aFNR) and CLPB (aCLPB) were used to assess their levels. Coomassie-stained gels indicate equal loading, and recombinant *A. thaliana* CLPB3 (rCLPB3) served as a positive control for aCLPB specificity. Numbers next to the blots represent molecular weight standards (in kDa). (B) Subcellular localization of CFP-tagged FNR WT and FNR Δ3 in *N. benthamiana* epidermal cells, visualized by confocal laser scanning microscopy. Fluorescence from CFP (cyan; i, v), chlorophyll autofluorescence (magenta; ii, vi), and brightfield views (iii, viii) are shown alongside merged images. Blue green and yellow circles highlight regions where CFP fluorescence overlaps with chlorophyll signal, indicating chloroplast localization of FNR WT and Δ3, respectively. A yellow arrow in panel x marks a stromule in a chloroplast of the FNR Δ3 variant. Scale bars: 20 μm.

One known molecular signature of the response to LIN treatment is the increase in the abundance of the chloroplastic chaperone CLPB (Llamas & Pulido, 2022). We have previously demonstrated the disaggregase activity of CLPB3 from *A. thaliana* (Parcerisa et al., 2020). We, therefore, investigated its abundance in whole leaf extracts and isolated plastids from agroinfiltrated leaves expressing FNR WT and FNR Δ3. At 48 h post-infiltration, a marked increase in CLPB levels was observed in leaves expressing FNR Δ3. In contrast, CLPB levels in FNR WT-expressing leaves were comparable to those of controls. CLPB levels further increased at 90 h, indicating a sustained response (Figure 2A). In isolated plastids from FNR Δ3-expressing leaves, the accumulation of CLPB was even more pronounced compared to FNR WT and control samples (Figure 2A).

Confirmation of the accumulation of FNR WT and, especially, FNR Δ3 in chloroplasts was obtained using confocal microscopy with variants fused to CFP. Signals were confined to chloroplasts, as judged by colocalization with chlorophyll autofluorescence (Figure 2C). Consistent with Western blot analyses, the CFP-FNR WT signal exhibited greater intensity and a broader distribution than the CFP-FNR Δ3 signal. Altogether, the data indicate that our chloroplast-targeted expression system can be used to trigger a CLPB-mediated unfolded protein response.

### A. thaliana plants expressing FNR deletion variants show a cpUPR

Having established that the expression of a misfolded protein in chloroplasts leads to increased levels of the disaggregase CLPB in transient expression assays, we next aimed to investigate the molecular features of this response in greater detail. To this end, we expanded our experimental model in two ways. First, we adopted a stable expression system in *A. thaliana*, which provides flexibility to explore developmental conditions, robustness, and simplicity. Second, we introduced an additional FNR folding variant more prone to aggregation than FNR Δ3. Specifically, the deletion of the C-terminal 20 amino acids in FNR results in a complete loss of solubility; this mutant, referred to as FNR Δ20, is unable to fold. Including this variant enabled us to probe how the degree of misfolding affects the chloroplast’s proteostasis machinery.

We transformed flowering *A. thaliana* plants using the floral dip method with *A. tumefaciens* strains harboring the same plasmids used in transient expression assays. An additional strain carrying pCHF3 with the coding sequence for TP-FNR Δ20 was included. The expression of the pea FNR proteins in *A. thaliana* T0 plants was assessed using orthogonal techniques. Immunodetection of the FNRs in *A. thaliana* leaves yielded similar results to those of *N. benthamiana*. Again, the anti-FNR antibodies cross-react between the endogenous *A. thaliana* FNR isoforms (FNR1 and FNR2) and the alien pea FNR isoforms. An increased signal was detected only in plants overexpressing FNR WT (Figure 3A). A double band was visible in these plants, consistent with the molecular mass difference between *A. thaliana* FNRs in their mature forms (35.3 kDa) and pea FNR WT (34.8 kDa). The folding-defective variants FNR Δ3 and FNR Δ20 are not present in sufficient quantities to be detected by this technique. So, the expression and chloroplast localization of FNR WT and the folding variants were assessed by confocal microscopy using CFP fusions. Results mirrored those in *N. benthamiana*: FNR WT–CFP showed strong organellar localization, while the folding variants exhibited weaker and less abundant CFP signals (Figure 3B).

**Figure 3:**
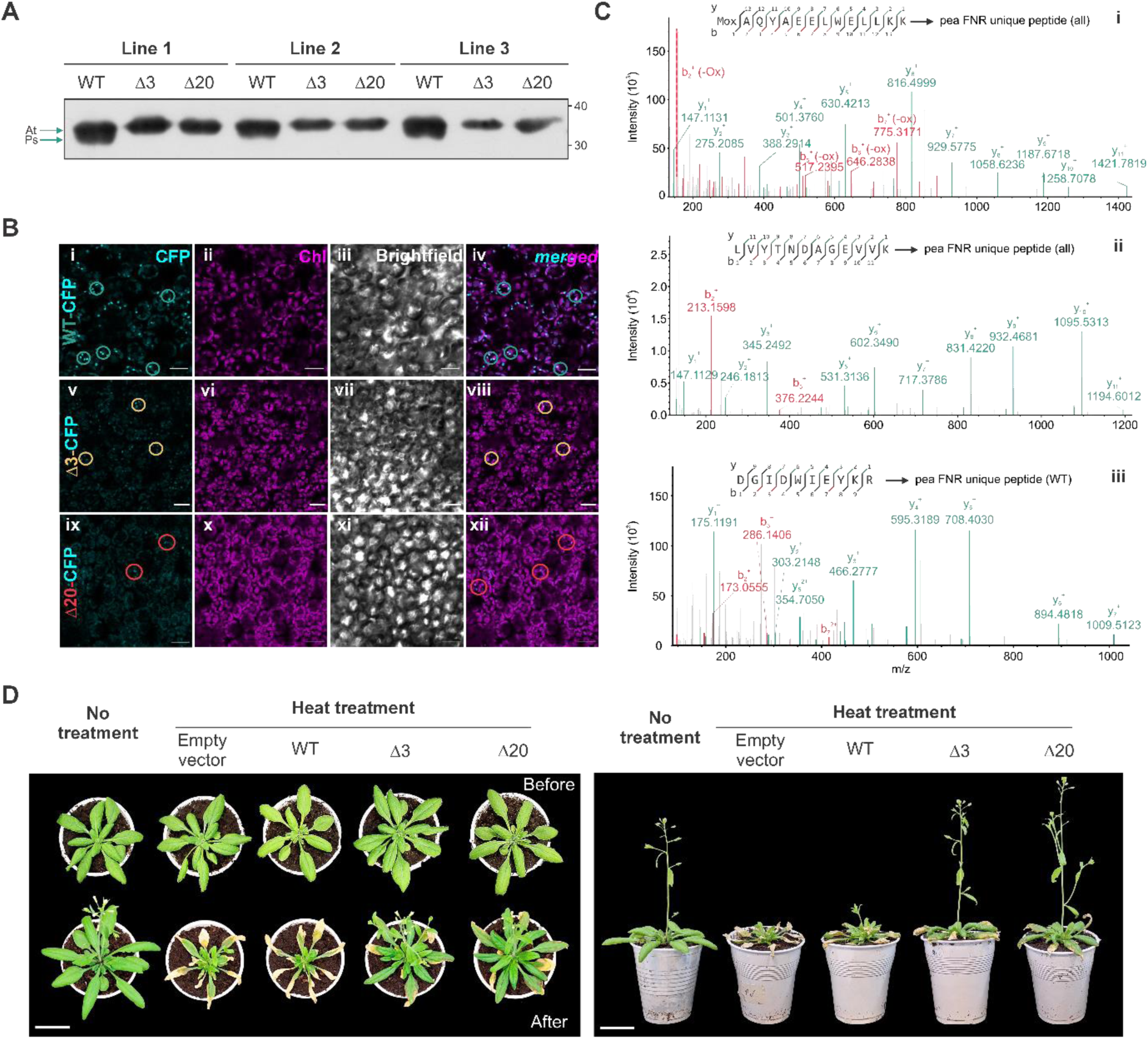
Expression, localization, and physiological impact of FNR variants in *A. thaliana*. (A) Immunoblots showing FNR levels in three representative and independent transgenic lines of *A. thaliana* expressing pea FNR WT, FNR Δ3, or FNR Δ20. Equal loading was confirmed by staining of total protein (bottom panel). The anti-FNR antibody (aFNR) detects endogenous *A. thaliana* isoforms (FNR1 and FNR2) and exogenous pea FNRs. A double band is observed in FNR WT-expressing plants, consistent with the size difference between endogenous and pea FNRs (arrows; At: *A. thaliana*, Ps: *Pisum sativum*. Numbers next to the blots represent molecular weight standards (in kDa). (B) Confocal laser scanning microscopy images of leaf epidermal cells from *A. thaliana* plants stably overexpressing CFP-FNR WT (i – iv), CFP-FNR Δ3 (v – viii), or CFP-FNR Δ20 (ix – xii). The CFP fluorescence signal (cyan) is shown in panels i, v, and ix; chlorophyll autofluorescence (magenta) is shown in panels ii, vi, and x; and corresponding brightfield images in panels iii, vii, and xi. Merged images (iv, viii, xii) combine CFP and chlorophyll signals to assess colocalization. Colored circles highlight representative regions where CFP signal overlaps with chlorophyll autofluorescence, indicating plastid localization: blue-green in i and iv (FNR WT), orange in v and viii (FNR Δ3), and red in ix and xii (FNR Δ20). Scale bars: 20 µm. (C) (i) Spectrum of the MAQYAEELWELLKK bearing and oxidized methionine residue (Mox), specific to all pea FNR variants (WT, Δ3, and Δ20). (ii) Spectrum of the variant-specific peptide LVYTNDAGEVVK, also shared by FNR WT, Δ3, and Δ20. (iii) Spectrum of the peptide DGIDWIEYKR, found exclusively in plants expressing pea FNR WT, as it contains the DWI motif absent in FNR Δ3 and FNR Δ20. b- and y-ion series are indicated in red and blue-green, respectively. (D) *Arabidopsis* plants expressing wild-type or mutant FNR variants (Δ3 and Δ20) were exposed to heat stress (45 °C) and evaluated for phenotypic responses. Top views (left) show representative plants before (top row) and 7 days after treatment (bottom row). Scale bar: 3.25 cm. Side views (right) reveal differences in bolting and overall growth. Scale bar: 3.5 cm.

To obtain more evidence that the pea FNR variants were being expressed in transformed *A. thaliana* plants, we sought to detect unique peptides of the pea variants in leaf extracts by mass spectrometry. Unique peptides refer to tryptic peptides that are only found in the pea variants and are absent in the endogenous FNRs. For example, the tryptic peptides “MAQYAEELWELLKK” and “LVYTNDAGEVVK” are specific to FNR WT, FNR Δ3, and FNR Δ20. These peptides were indeed detected in leaf extracts from all plants expressing pea FNRs, indicating their successful expression (Figure 3C, i and ii). Of note, the pea FNR-specific tryptic peptide “DGIDWIEYKR” (which includes the DWI triad removed in FNR Δ3 and from where the deletion in FNR Δ20 starts) could only be found in plants expressing pea FNR WT (Figure 3C, iii). Consistently, the number of peptide spectrum matches of pea FNR peptides in plants expressing FNR WT was higher than in plants expressing the mutants. This again reflects its greater abundance in leaves. Finally, we could not detect any tryptic peptide from the TP, which suggests that preproteins were being readily processed into their mature form.

Despite the overexpression of FNR WT and the folding-defective variants, all transgenic lines showed normal growth and development under standard conditions throughout generations, with no evident phenotypic alterations. To assess whether this apparent lack of phenotype persisted under proteotoxic stress, we exposed transgenic and control seedlings to elevated temperature, a condition known to challenge protein homeostasis. Remarkably, plants expressing FNR Δ3 and Δ20 displayed greater resilience to heat stress (45 °C, 12 h) than wild-type or FNR WT-expressing controls, maintaining greener rosettes and sustained post-treatment growth (Figure 3D). Altogether, these observations suggested a primed quality control system in cpUPR-activated lines.

### The degree of misfolding shapes the cpUPR response

To investigate whether the observed heat protection was associated with cpUPR activation, we analyzed the leaf proteome using high-resolution mass spectrometry. Given the different solubility and presumed folding states of FNR Δ3 and Δ20, we hypothesized that the cpUPR would scale with the severity of protein misfolding. Protein extracts from four biological replicates of *A. thaliana* expressing pea FNR WT, Δ3, or Δ20 were subjected to tryptic digestion and analyzed via a label-free quantification strategy. Intragroup Pearson correlation coefficients were above 0.96, indicating excellent reproducibility and technical consistency (Supplemental Figure 1A). Principal component analysis revealed a clear separation between samples expressing FNR WT and those expressing the folding-defective variants, suggesting a distinctive change in the proteome under these conditions (Supplemental Figure 1B).

Next, we analyzed the differential abundance of proteins that were significantly regulated. The expression of FNR Δ3 resulted in an increased abundance of 156 proteins and a downregulation of 26, compared to FNR WT. At the same time, FNR Δ20 variant induced a broader shift, with 239 proteins upregulated and 17 downregulated, suggesting a more extensive response to this aggregation-prone variant (Supplemental Table 1). Figure 4A summarizes the number of differentially expressed proteins and illustrates the overlaps and specificities among the different comparisons. Notably, over 90% of the DEPs observed in Δ3-expressing plants were also present in the Δ20 background, indicating that the response to FNR Δ3 represents a subset of the more extensive alterations induced by Δ20. To visualize expression patterns, we generated a heatmap of all differentially expressed proteins across genotypes (Figure 4B). Hierarchical clustering separated the genotypes and identified two prominent protein groups. One cluster is upregulated only in Δ20, while the other shows upregulation from WT to Δ3 and Δ20. These trends suggest the activation of both shared and variant-specific pathways.

**Figure 4.**
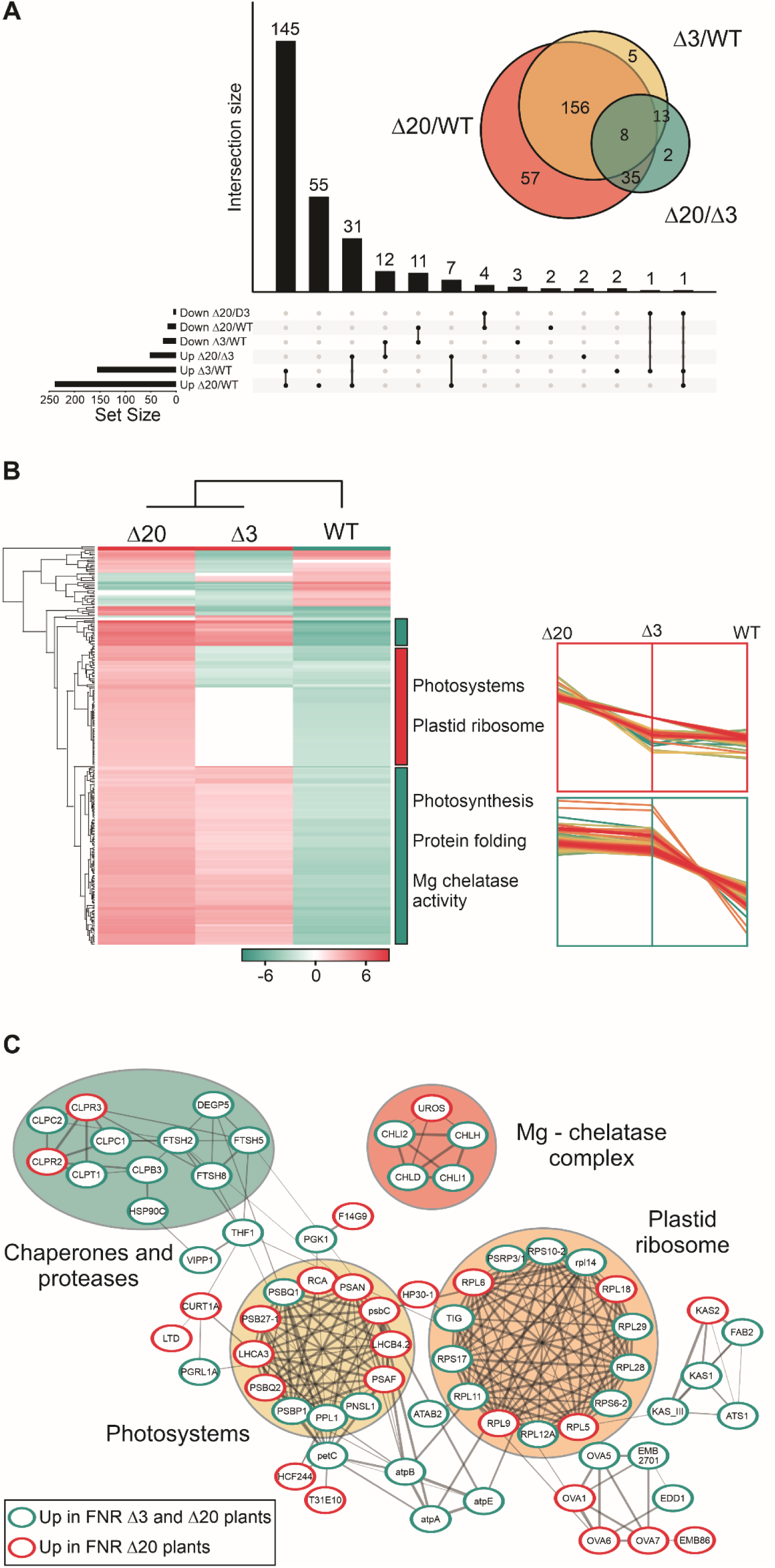
Quantitative proteomics reveals cpUPR activation and functional reprogramming in response to misfolded FNR variants. (A) UpSet plot with Venn diagram inset showing the number and overlap of differentially expressed proteins in Δ3/WT, Δ20/WT, and Δ20/Δ3 comparisons. Bars represent the size of intersections between sets, with up- and downregulated proteins indicated. The Venn diagram summarizes the overlap of differentially expressed proteins among the three comparisons. (B) Heatmap of significantly regulated proteins across genotypes (Δ20, Δ3, and WT), based on hierarchical clustering. The color gradient below the heatmap represents the values of the Z-score presented in Supplemental Table 1. Bars to the right of the plot encompass enriched terms for proteins upregulated in plants expressing FNR Δ3 and FNR Δ20 (blue-green bars) or FNR Δ20 (red bar). Line plots (right) illustrate expression trends for selected clusters, indicating both shared and variant-specific proteomic responses. Two major clusters are depicted: one comprising proteins upregulated only in FNR Δ20-expressing plants (red marquee) and another showing an increase in abundance from FNR WT to FNR Δ3 and Δ20 (blue-green marquee) (C) These clusters were used to generate a protein-protein interaction network of chloroplastic proteins upregulated in Δ3 and/or Δ20 lines (STRING confidence ≥ 0.7). Nodes represent individual proteins, with red borders indicating proteins uniquely upregulated in the FNR Δ20 background and blue-green borders denoting proteins upregulated in both FNR Δ3 and Δ20 genotypes.

Biological interpretation of the proteomics data was aided by network and enrichment analysis. We focused on proteins with significantly increased levels in plants expressing FNR Δ20 and Δ3 compared to controls expressing FNR WT (152 proteins) or those with increased levels only in the FNR Δ20 lines (87 proteins). The Uniprot accession codes for these 239 proteins were used to interrogate the STRING database at a 0.7 confidence threshold. The resulting network was densely connected and presented a few singletons. Enrichment analysis revealed that the term “chloroplast” was significantly enriched, as 50% of the queried proteins were chloroplastic, indicating that half of the proteome variation occurs in this organelle. Regarding the molecular function of these selected chloroplastic proteins, they are grouped into clusters under the terms “chaperones and proteases,” “plastid ribosome,” and “photosystems,” among others. Chloroplastic chaperones, proteases, and members of the plastid ribosome were detected to have altered levels in response to FNR Δ3 and FNR Δ20. On the other hand, many photosynthesis-associated proteins showed increased levels predominantly in response to FNR Δ20 (Figure 4B and C).

Our quantitative proteomics analysis allowed us to measure the levels of key chloroplastic chaperones involved in plastidic protein homeostasis. CLPB3 and chloroplastic HSP90C (HSP90-5) displayed increased abundance in both FNR Δ3 and Δ20 overexpressing lines. In contrast, the levels of chloroplastic HSP70s remained unchanged (HSP70.1 and HSP70.2) (Figure 5A). The CLP system was also upregulated both in the chaperone complex subunits CLPC1 and CLPC2, as well as with the components of the proteolytic core CLPR2 and CLPR3 and the accessory assembly subunit CLPT1 (Supplemental Table 1). We confirmed the levels of some of the candidate proteins involved in the cpUPR by Western blot. The levels of CLPB3, the chloroplastic disaggregase from *A. thaliana*, as well as those of CLPC (CLPC1 and CLPC2), significantly increased by as much as six (CLPB3) and three (CLPC) times in plants expressing FNR Δ3 or FNR Δ20 compared to controls or plants expressing FNR WT (Figure 5B). As previously observed for CLPB in *N. benthamiana*, the expression of aggregation-prone FNR variants in *A. thaliana* chloroplasts leads to an increase in the levels of quality control proteins to sustain proteostasis. Altogether, the analysis of the expression of these central chaperones and proteases indicates precise upregulation of the chloroplast proteostasis machinery in response to the expression of FNR Δ3 and Δ20.

**Figure 5.**
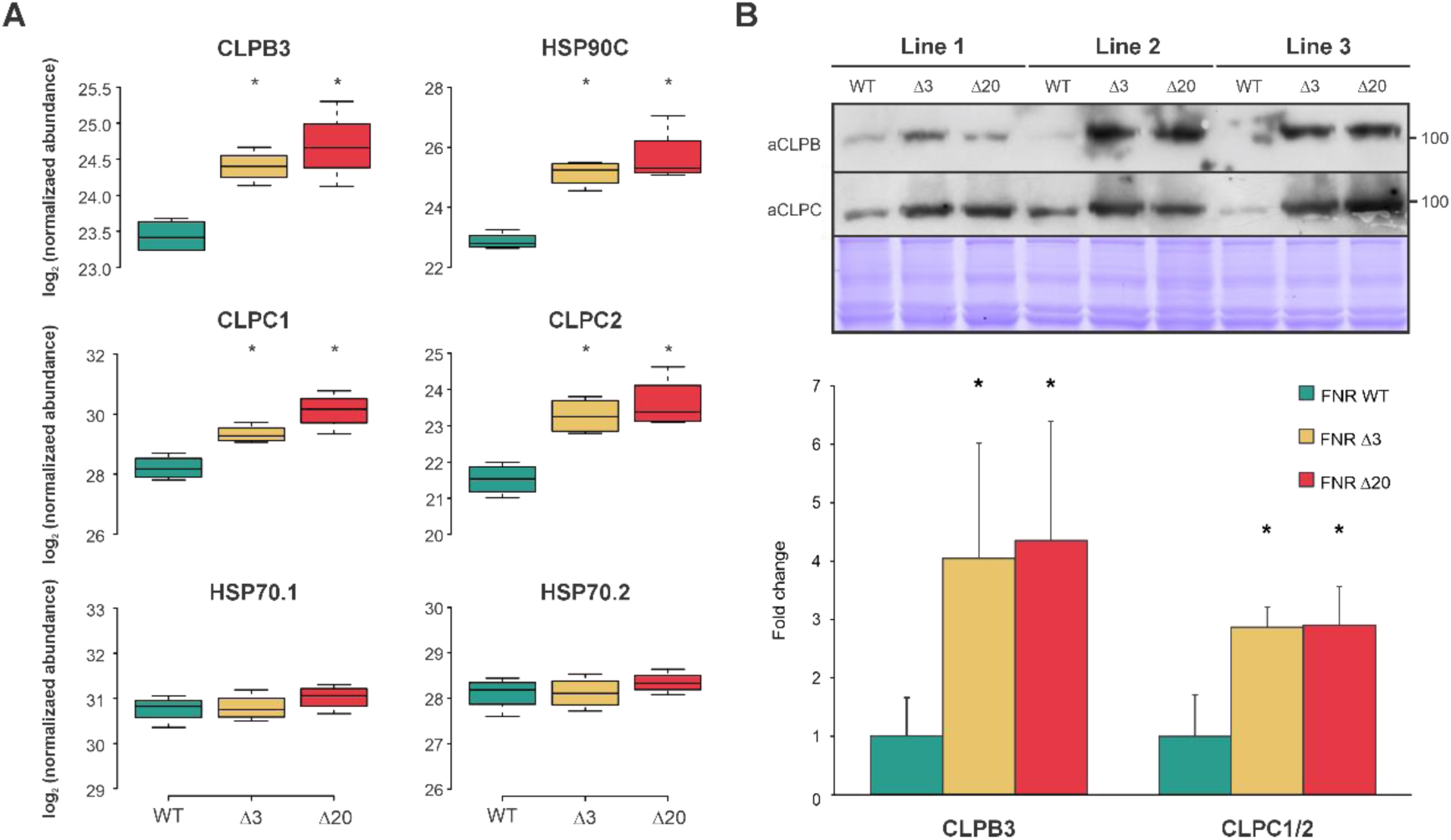
Abundance of selected chloroplastic chaperones in FNR overexpressing lines. (A) Boxplots showing log2-transformed normalized abundances of CLPB3, CLPC1, CLPC2, HSP90C, HSP70.1, and HSP70.2 in FNR WT, Δ3, and Δ20 backgrounds. Asterisks indicate statistically significant differences (p < 0.05) compared to the wild type (WT). (B) Immunoblots showing CLPB3 and CLPC1/2 levels in three representative and independent transgenic lines of *A. thaliana* expressing pea FNR WT, FNR Δ3, or FNR Δ20. Equal loading was confirmed by staining of total protein (bottom panel). aCLPB and aCLPC, antibodies against CLPB3 and CLPC (both CLPC1 and 2), respectively. Quantification of protein level is shown in the bar plot below the immunoblots. Bars represent mean fold change ± SD. Asterisks denote statistically significant differences relative to FNR WT (p < 0.05).

### The proteomic signature of the cpUPR differs from that of the chemically induced UPR

A large body of evidence for a cpUPR stems from research using plants grown in the presence of LIN and mutants in genes encoding components of the protein quality control (PQC) system (Gao et al., 2023; Llamas & Pulido, 2022). LIN has a dose-dependent effect, ranging from a blockage of plastid expression and accumulation of protein aggregates to a proteotoxic effect as a consequence of perturbation of the import and folding of photosynthetic proteins. In particular, concentrations as high as 550 μm are used to trigger plastidic retrograde signals (Di Silvestre et al., 2025; Koussevitzky et al., 2007; X. Zhao et al., 2018). To discriminate between specific responses caused by the buildup of aggregated proteins (as in *Arabidopsis* lines overexpressing FNR variants) and the broader effect of LIN treatment, we compared these two approaches at a proteomic level. To minimize the deleterious effects of LIN on photosynthetic performance and using ClpB3 levels as a proxy for the imbalance in proteostasis, we selected 15 μM LIN. Under this condition, CLPB3 levels were more than three times higher compared to the controls (Supplemental Figure 2A). At the same time, the plants exhibited little phenotypic defect, i.e., low chlorosis and smaller size (Supplemental Figure 2B).

Quality control of the proteomics data indicated an excellent correlation between samples and separation of the groups in PCA plots (Supplemental Figure 3), suggesting changes in the leaf proteome caused by exposure to 15 μm LIN.

However, the set of proteins with altered abundance in seedlings treated with 15 μM LIN differed markedly from those observed in plants expressing the aggregation-prone FNR variants. Among the chloroplast-localized quality control proteins, only CLPB3 and FTSH8 showed an increase in abundance approaching twofold. In contrast, most chaperones and proteases upregulated in the FNR variant-expressing plants showed minimal changes or fell below the significance threshold in LIN-treated samples (Figure 6).

**Figure 6.**
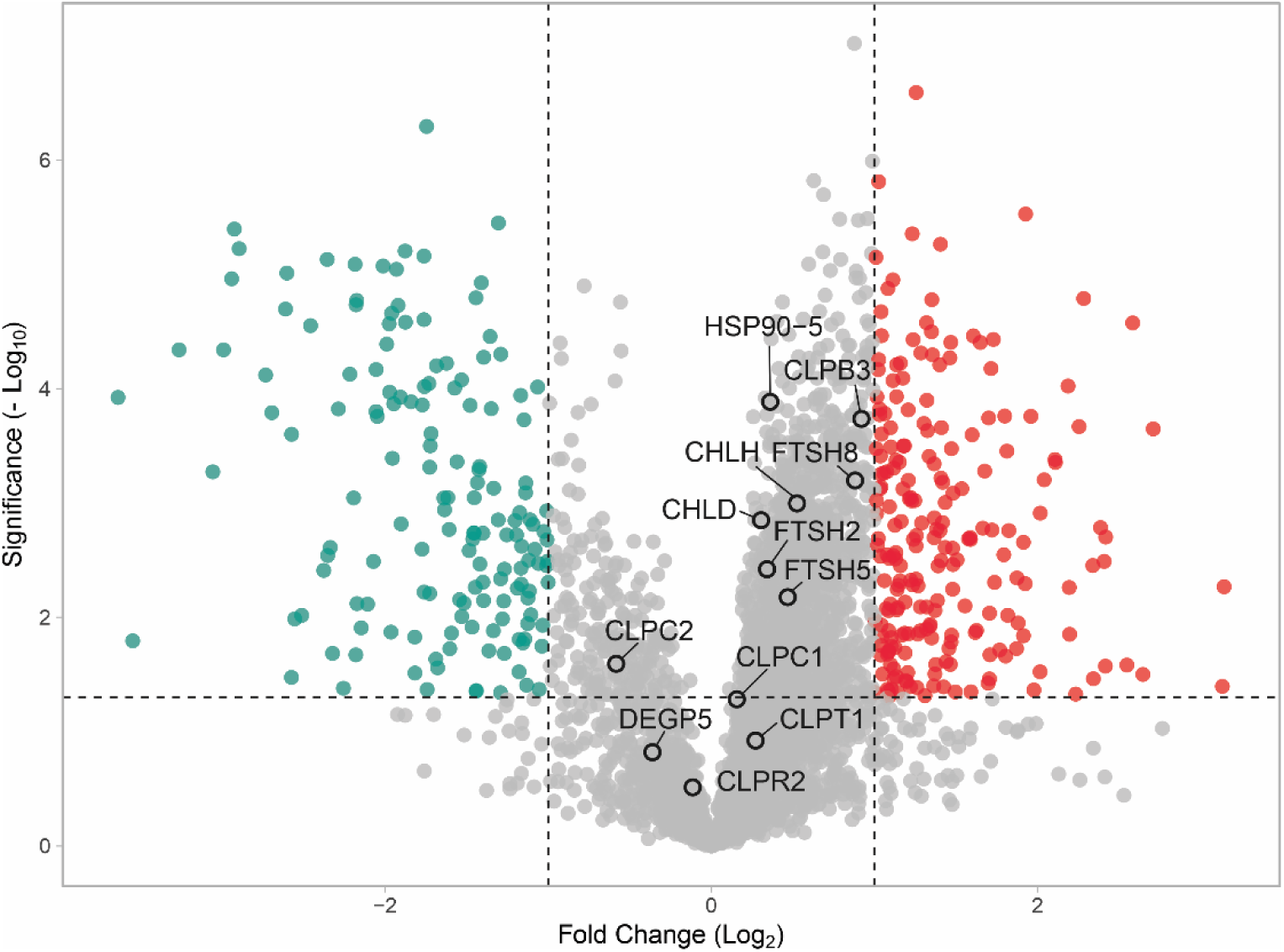
Differential protein abundance in LIN-treated *A. thaliana* seedlings. Volcano plot showing changes in protein abundance in 14-day-old Col-0 seedlings treated with 15 μM lincomycin (LIN) compared to untreated controls. Proteins significantly upregulated (red) or downregulated (blue-green) are shown, based on thresholds of (log₂ fold change) > 1 and adjusted p < 0.05 (dashed lines). Highlighted proteins include chloroplast-localized chaperones and proteases.

Proteins with significantly elevated levels were loosely connected, and singletons were abundant. There were no enriched Gene Ontology terms for this group. Proteins with considerably lower levels formed an extensive network comprising two major clusters: ribosomal proteins and proteins involved in photosynthesis (Supplemental Figure 4 and Supplemental Table 2). The overall comparison between FNR variants- and LIN-triggered UPR reveals two types of response. LIN-treated plants showed a slight increase in levels of PQC members while broadly repressing plastid gene expression and downregulation of photosynthetic proteins. In contrast, the response to the presence of FNR Δ3 and FNR Δ20 contributed differently to restoring proteostasis by inducing a more complex set of PQC components and enhancing plastid gene expression, ultimately improving chloroplast functionality under this condition.

Our findings support a model in which chloroplasts can mount a specific cpUPR to adapt to the accumulation of misfolded proteins, restoring proteostasis without compromising organelle function (Figure 7). In contrast to the LIN-triggered cpUPR-like response characterized by inhibition of plastid gene expression, a partial induction of PQC components, and chronic stress and chlorosis, the overexpression of folding-defective FNR variants (Δ3 and Δ20) activates a cpUPR that restores proteostasis. The cpUPR involves retrograde signaling to the nucleus and subsequent induction of nuclear genes encoding for PQC components, including CLPB3, CLPC, HSP90.5, and FTSH and DEG proteases, along with photosynthesis-associated nuclear genes (PhANGs). This response appears well tolerated: no visible phenotypes or developmental arrests were observed, and cpUPR activation conferred protection against heat stress. Therefore, our results revealed a canonical cpUPR capable of sustaining chloroplast functionality.

**Figure 7.**
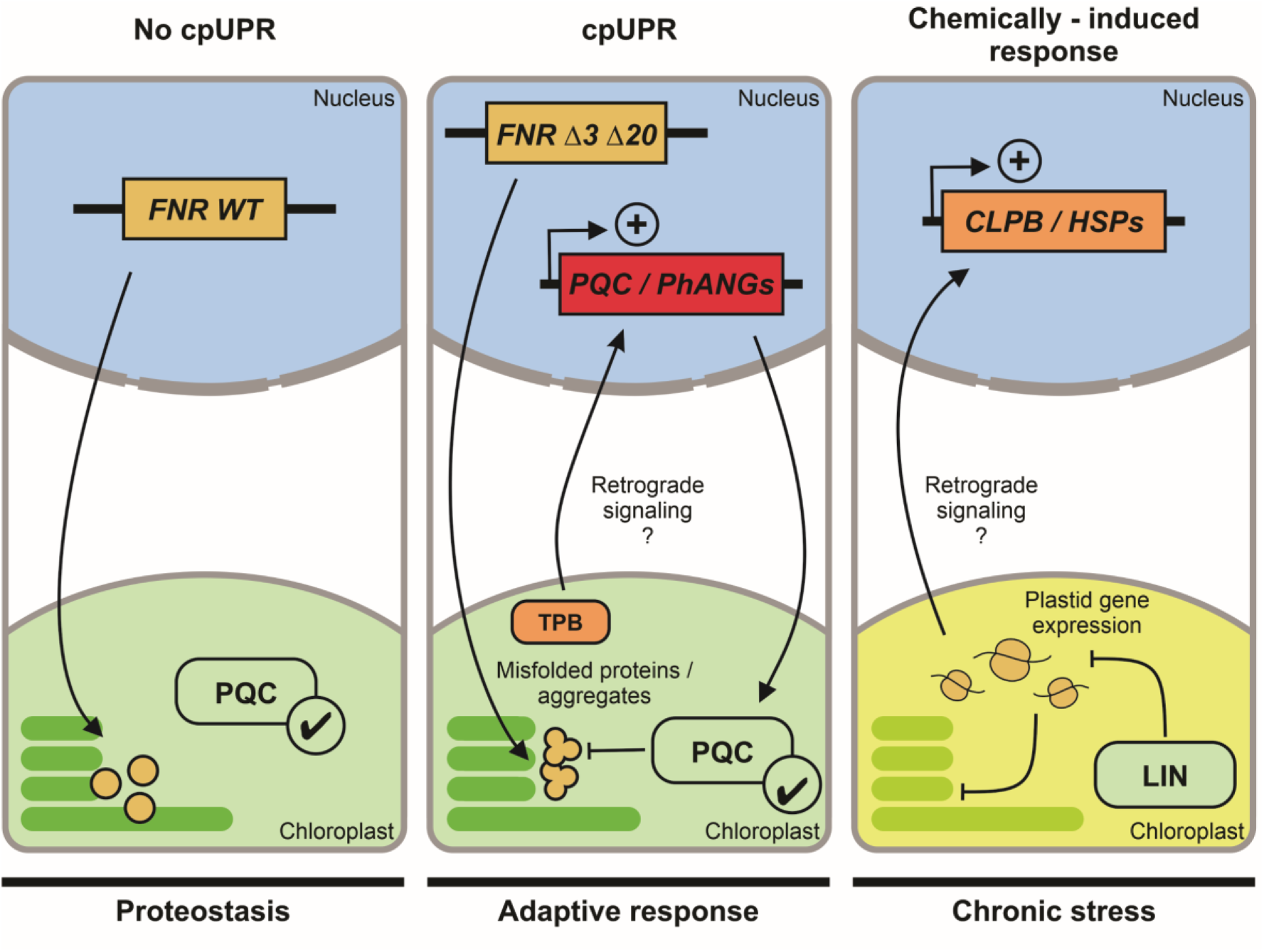
A model for cpUPR activation by folding-defective proteins versus chemical stress. In the absence of chloroplast stress (left panel), overexpression of FNR WT does not impair proteostasis, and the PQC system operates at basal levels. Overexpression of folding-defective FNR variants (Δ3 or Δ20, middle panel) results in aggregation, triggering retrograde signaling, possibly mediated by the tetrapyrrole biosynthesis (TPB) pathway, and upregulation of nuclear genes encoding PQC components (e.g., CLPB3, CLPC, HSP90.5, FTSH, DEG) and photosynthesis-associated nuclear genes. This cpUPR activation is specific and well tolerated, with no visible phenotypes and improved thermotolerance. In contrast, LIN treatment (right panel) inhibits plastid gene expression and induces a cpUPR-like response; however, persistent translational arrest leads to unresolved stress, impaired development, and chlorosis.

## DISCUSSION

Over the last decade, numerous lines of research have pointed out an unfolded protein response in chloroplasts using several strategies like mutants in PQC components and LIN treatments (Di Silvestre et al., 2025; Dogra et al., 2019; Llamas et al., 2017; Ramundo & Rochaix, 2014a and reviewed in Gao et al., 2023; Llamas & Pulido, 2022). To add definitive evidence for its existence, we drew inspiration from earlier work in mitochondria and the ER, where the buildup of aggregation-prone proteins unleashed the corresponding UPR. In this way, the overlap of responses induced by global triggers (such as chemicals and abiotic stress) is diminished, and the molecular actors of the UPR can thus be uncovered. We chose variants of FNR that our group designed and characterized more than 30 years ago (Orellano et al., 1993). In these, three or twenty residues were eliminated from the C-terminal end, preventing proper folding (FNR Δ3 or Δ20, respectively). As evidenced by mass spectrometry identification and fluorescence microscopy, the reductases were expressed in leaves and accumulated in chloroplasts. This was expected, at least, for the wild-type protein, as transgenic tobacco plants overexpressing pea FNR had already been reported. As described by Rodriguez et al., the overexpression of pea FNR does not affect growth or photosynthetic parameters under normal conditions (Rodriguez et al., 2007). However, the accumulation of FNR WT was several times higher than that of the folding defective variants, even though all genetic constructs used for plant transformation were the same. This suggests prompt degradation of the misfolded variants as part of a functional proteostatic response, in line with the increase in protease content of the chloroplast.

The induction of *A. thaliana* CLPB3 and CLPC in the presence of the FNRs variants was part of a more complex response to restore proteostasis. Proteomic analysis revealed a coordinated increase of chaperones, proteases, and accessory assembly factors of the PQC system in response to FNR Δ3 and Δ20, most of them targeted to chloroplasts. FTSH is an ATP-dependent zinc metalloprotease complex anchored in the thylakoid membrane, comprised of four major isoforms (1, 2, 5, and 8) (Van Wijk, 2024). Higher amounts of FTSH 2, 5, and 8 were detected in FNR Δ3 and Δ20 overexpressing plants compared to overexpression of wild-type FNR. Another induced plastidic protease was DEG5, belonging to the DEG protease (formerly DEGP), which are ATP-independent Ser-type endopeptidases in the thylakoid lumen and stroma. These protease complexes are central for chloroplast proteostasis, and they act together to degrade photodamaged D1 subunits of the PSII reaction center (reviewed in (Yoshioka-Nishimura & Yamamoto, 2014). We also observed a substantial accumulation of components of the stromal CLP protease. None of these proteases usually degrade appropriately folded proteins. Therefore, they are presented with substrates that adaptors and molecular chaperones have previously recognized (Aguilar Lucero et al., 2021; Rosano et al., 2012). For example, CLPC1 and C2 are HSP100 hexameric chaperones that unfold damaged proteins and translocate them through their central pore to the barrel-shaped CLPP protease (Bruch et al., 2012; Rosano et al., 2011). Levels of both CLPC1 and C2 were increased in *A. thaliana* plants overexpressing the folding-defective FNR variants. The abundance of the assembly protein CLPT1 (Colombo et al., 2014; Kim et al., 2015) and the proteolytic subunits CLPR2 and CLPR3 increased as well (Kim et al., 2009). The ultimate fate of a misfolded protein is not always degradation; a subset of chaperones also rescues them from aggregates and partially folded intermediates. The major chloroplastic disaggregase is CLPB3, which can refold aggregated proteins *in vitro* (Parcerisa et al., 2020). HSP90.5 is vital for chloroplast biogenesis and embryogenesis, while its interaction partner VIPP1 is essential for the biogenesis and maintenance of thylakoid membranes (Feng et al., 2014). These three proteins showed higher levels in response to FNR Δ3 and Δ20, underscoring the molecular response to their presence. Based on these results, our proteomic analysis of the leaf proteome in the presence of FNR Δ3 and Δ20 provides a *bona fide* description of the PQC as part of the activated cpUPR to sustain proteostasis.

Most components of the PQC system are encoded in the nucleus, implying that their induction relies on retrograde signals from the organelle to the nucleus to restore proteostasis (Barajas-López et al., 2013; Gao et al., 2023). Mounting evidence highlights GUN1 as an important regulator of chloroplast stress responses (Gao et al., 2023; Llamas & Pulido, 2022; Wu & Bock, 2021). GUN1 is a pentatricopeptide repeat-containing protein that integrates multiple retrograde signaling pathways (Wu & Bock, 2021). During early plant development, it represses expression of PhANGs in response to plastid dysfunction, particularly upon treatment with inhibitors such as LIN or norfluorazon, the latter triggering plastid gene expression pathways through photooxidative damage (Koussevitzky et al., 2007; Wu et al., 2018; Wu, Meyer, Wu, et al., 2019; X. Zhao et al., 2018). Mutants impaired in plastid gene expression or PQC components often display plastid-dependent phenotypes (e.g., pale or variegated leaves) that are suppressed by loss of GUN1 function, further implicating GUN1 in stress-related retrograde signaling (Gao et al., 2023; Marino et al., 2019; Tadini et al., 2016, 2020; Wu, Meyer, Wu, et al., 2019).

In our proteomic analysis, GUN1 was not detected in plants expressing FNR Δ3, Δ20, or treated with LIN. Although this prevents concluding its role in the cpUPR, several factors may explain its absence. First, our experiments were performed in 14-day-old *Arabidopsis* seedlings, a stage when GUN1 is post-transcriptionally repressed (Wu et al., 2018). Second, the strong induction of CLPC1 and CLPC2 in response to the FNR variants may promote GUN1 degradation, as previously reported (Wu et al., 2018). Third, both GUN1 levels and the severity of the *gun* phenotype depend on the intensity and timing of chloroplast stress (Llamas et al., 2017; Tadini et al., 2020; Wu et al., 2018; X. Zhao et al., 2018). The mild LIN treatment used here (15 μM) likely contributes to the lack of GUN1 detection yet still impairs chloroplast function by suppressing photosynthetic gene expression and protein import. These detrimental effects have recently been shown to extend to the cytosol, where the overaccumulation of chloroplast precursor proteins triggers a heat-shock–like response and transcriptional reprogramming (Hong et al., 2025). In contrast, in our FNR variants model, the degree of proteotoxic effect is so gentle that the cpUPR can be activated and studied in phenotypically normal seedlings, offering a more nuanced view of the organelle’s ability to communicate and recover from the imbalance in proteostasis.

Interestingly, together with the general induction of photosynthetic genes (see below), we observed an increase in the enzymes of the tetrapyrrole biosynthesis pathway (TPB). Specifically, all subunits of the Mg-chelatase complex (*i.e.*, CHLI1 and 2, CHLD, and CHLH) showed increased levels in plants overexpressing the folding-defective FNRs. The complex plays a fundamental role in chlorophyll biosynthesis by catalyzing the insertion of a magnesium ion into protoporphyrin IX (Proto IX). This commits the precursor to chlorophyll biosynthesis, a pathway essential for producing the pigment, at the expense of the heme branch, which involves the insertion of an iron ion instead. Beyond its enzymatic role, the Mg-chelatase complex subunits (i.e., CHLH/GUN5 and its activator GUN4), as well as the branched pathway downstream of Proto IX (including HEME OXYGENASE1/GUN2, PHYTOCHROMOBILIN SYNTHASE/GUN3, and FERROCHELATASE1/GUN6), have been identified as sources of plastid-to-nucleus retrograde signals in the initial gun screening (Larkin, 2016; Mochizuki et al., 2001; Woodson et al., 2011). Although the nature of the intermediate steps remains under debate (Kindgren et al., 2012; Moulin et al., 2008; Strand et al., 2003), our results suggest that increased TPB activity is part of the cpUPR-associated gene expression, consistent with evidence from genetic and LIN-triggered responses (Hong et al., 2025; Wang et al., 2023).

In our work, the cpUPR was triggered by the expression of folding-defective proteins targeted to chloroplasts, drawing inspiration from the work of Zhao et al. that used a truncated version of OTC to unleash the mitochondrial UPR (Q. Zhao et al., 2002). Moreover, we used two variants with different solubility, permitting us to unveil the molecular players of the cpUPR as the protein crisis gets more severe. Overexpressing FNR Δ3 and Δ20 increases chloroplastic chaperones and proteases. However, FNR Δ20 adds a new layer of complexity, comprising proteins involved in membrane remodeling and further protection against stress. For example, SLOW GREEN 1 (SG1) is required for the early stage of chloroplast development. It may be involved in chloroplast protein biosynthesis and/or degradation (Hu et al., 2014). Photosystem II repair protein PSB27-H1 is engaged in the repair of photodamaged photosystem II (Nowaczyk et al., 2006). Protein CURVATURE THYLAKOID 1A determines thylakoid architecture by inducing membrane curvature (Armbruster et al., 2013). Protein HIGH CHLOROPHYLL FLUORESCENCE PHENOTYPE 244 is required for the biogenesis of photosystem II, especially for synthesizing the reaction center proteins (e.g., D1) (Li et al., 2019). Considering the functions of the proteins, we can speculate that the cpUPR is a mechanism to protect photosynthesis. As photosynthetic proteins are damage-prone, the organelle interprets the presence of aggregates as a general signal of injury, particularly damning to the photosynthetic electron transport chain.

Our comparative proteomic analysis of the stress response triggered by overexpressing the FNR variants and LIN shows that, while these processes share certain features, they also exhibit significant differences. CLPB3 and HSP90-5 showed increased levels under both treatments (although both were slightly below our chosen two-fold threshold in the proteomic analysis of plants under LIN treatment), which agrees with other reports using 15 μM LIN (Llamas et al., 2017). However, there was little overlap in the molecular response to low concentrations of LIN compared to our strategy. LIN treatment negatively impacted photosynthetic proteins and the plastid ribosome. Yet, its effect on chaperones, proteases, and other members of the PQC network was low. Strikingly, the levels of photosynthetic proteins showed opposite trends. Under LIN treatment, their abundance lowered, but a significant increase was detected in plants overexpressing the FNR variants. The former effect was expected as LIN causes chlorosis and disruption of the photosynthetic electron transport chain, leading to the degradation of its constituents. Building upon our hypothesis that the accumulation of the FNR variants is sensed as a mild decrease in photosynthetic performance induced by high light or heat stress (which mainly affects the electron transport chain), the upregulation of photosynthetic proteins may be a mechanism for their rapid replenishment to ensure proteostasis. The hypothesis of an acclimation mechanism triggered by FNR variants to keep chloroplasts functional is supported by a general and extensive induction of the plastidic gene expression machinery, a phenomenon that also occurs upon overaccumulation of stromal proteins (Tadini et al., 2023).

Our results also suggest that cpUPR activation may serve as an effective strategy to enhance thermotolerance in plants. Unlike conventional approaches that rely on overexpressing individual chaperones or proteases (e.g., CLPB3 or members of the HSP100 family), our system triggers a broad, coordinated proteostasis response by expressing folding-defective chloroplastic proteins without compromising cell health. This leads to the simultaneous upregulation of multiple quality control components, like disaggregases, chaperones, proteases, and thylakoid maintenance factors that likely act in concert to mitigate proteotoxic damage. As such, expressing a tailored misfolded protein or polypeptides targeted to specific organelles is a promising approach to boost plant resilience under challenging conditions such as heat stress. This concept positions the cpUPR as a tunable, systems-level approach with potential applications in crop improvement.

## ACKNOWLEDGMENTS

We thank Dr. Enrique Morales for technical support in confocal microscopy. This work was supported with a grant from the Agencia Nacional de Promoción de la Investigación, el Desarrollo Tecnológico y la Innovación (PICT 2019-02971) awarded to GLR.

