## Supplemental Figure 1 for "A canonical chloroplast unfolded protein response triggered by misfolded polypeptides"

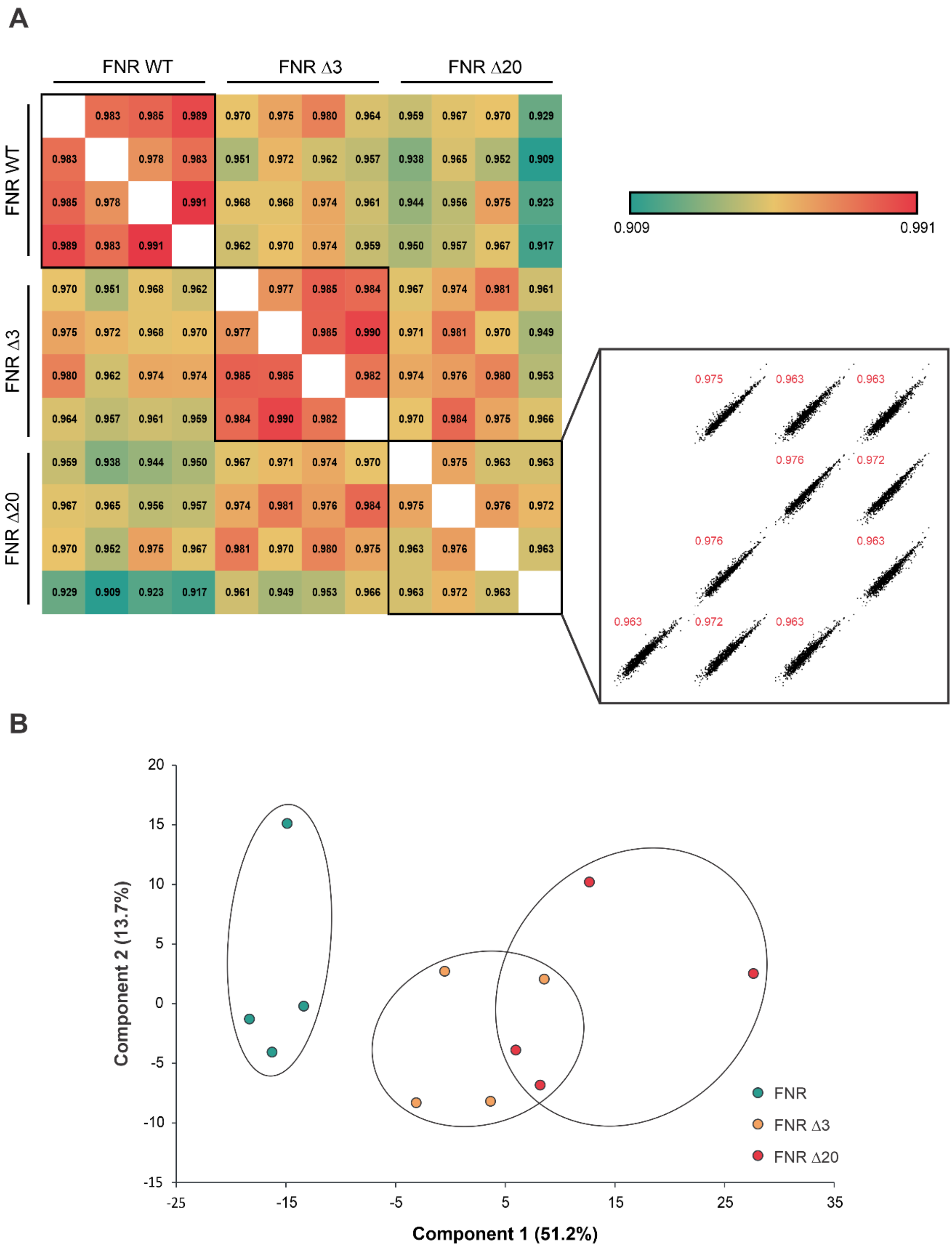


**Supplemental Figure 1: Proteomic analysis of *A. thaliana* plants expressing FNR variants.** (A) Pearson correlation matrix of label-free quantitative proteomics data from *A. thaliana* expressing pea FNR WT, FNR Δ3, or FNR Δ20. Four biological replicates were analyzed per condition. Inset: scatter plots of selected replicate comparisons, with correlation values indicated in red. (B) Principal component analysis of the proteomic profiles. Component 1 (51.2%) and 2 (13.7%) capture the major variance in the dataset. PCA reveals the separation of FNR WT samples from those expressing the folding-defective variants (FNR Δ3 and FNR Δ20), indicating distinct proteomic responses associated with the accumulation of misfolded FNR.

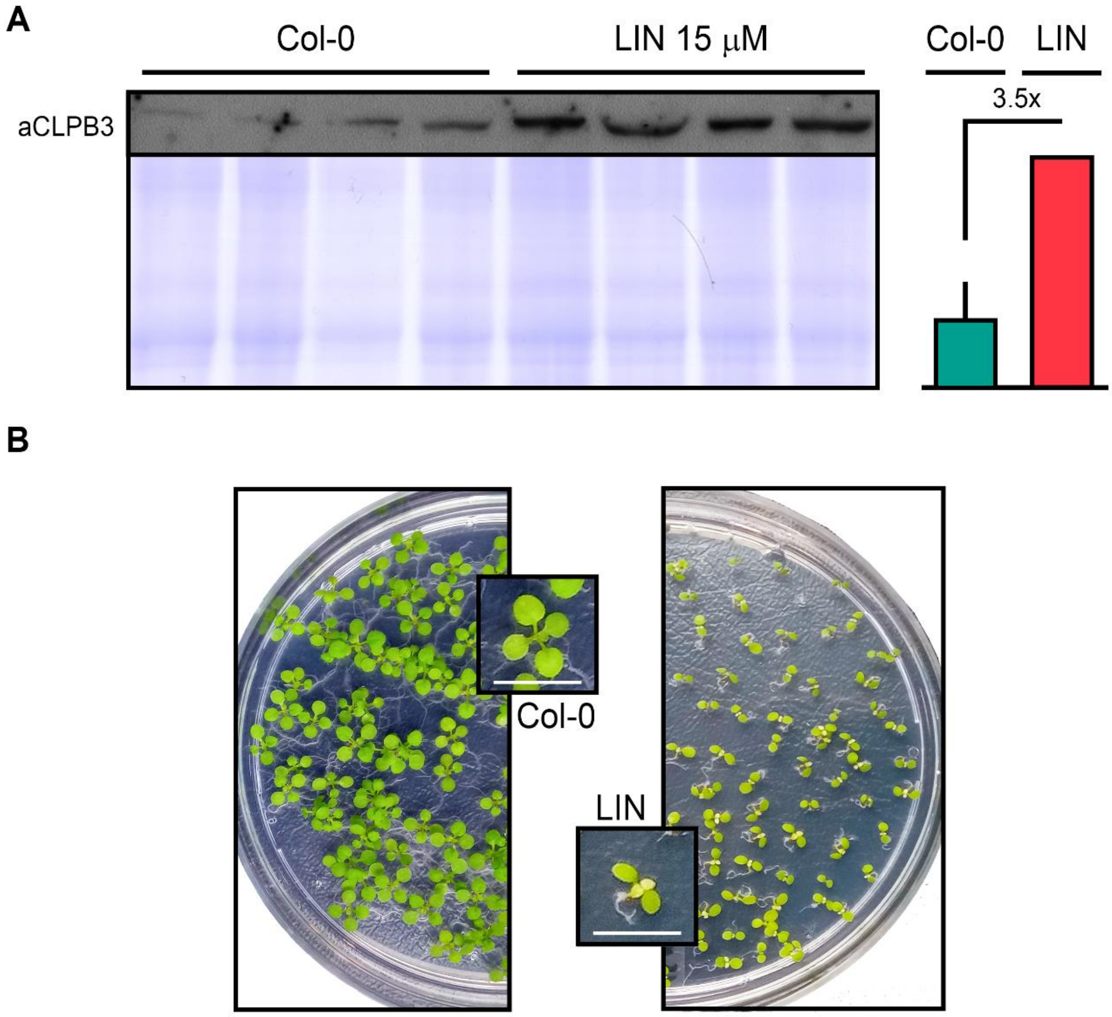


**Supplemental Figure 2: LIN induces CLPB3 accumulation and some phenotypic alterations in *A. thaliana* seedlings.** (A) Immunoblot analysis of CLPB3 levels in 14-day-old *A. thaliana* Col-0 seedlings treated with 15 µM lincomycin (LIN). Total protein extracts were probed with anti-CLPB3 antibodies (top panel), and Coomassie Brilliant Blue staining (bottom panel) was used as a loading control. Bars represent mean ± SD. (B) Representative images showing the phenotypic effects of LIN treatment on *A. thaliana* seedlings. Insets show close-up views of typical individuals from each condition. Scale bars = 5 mm.

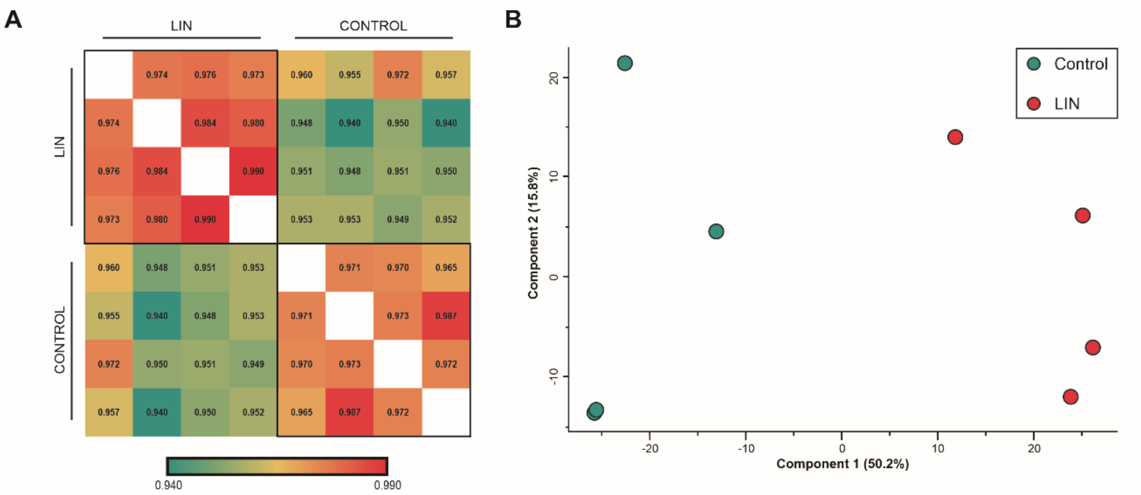


**Supplemental Figure 3. Quality control of proteomics data from LIN-treated and control samples.** (A) Pearson correlation heatmap of all biological replicates from LIN-treated and control samples. (B) Principal component analysis of the same datasets shows a clear separation between LIN-treated (red) and control (blue-green) samples along the first principal component (PC1, 50.2%).

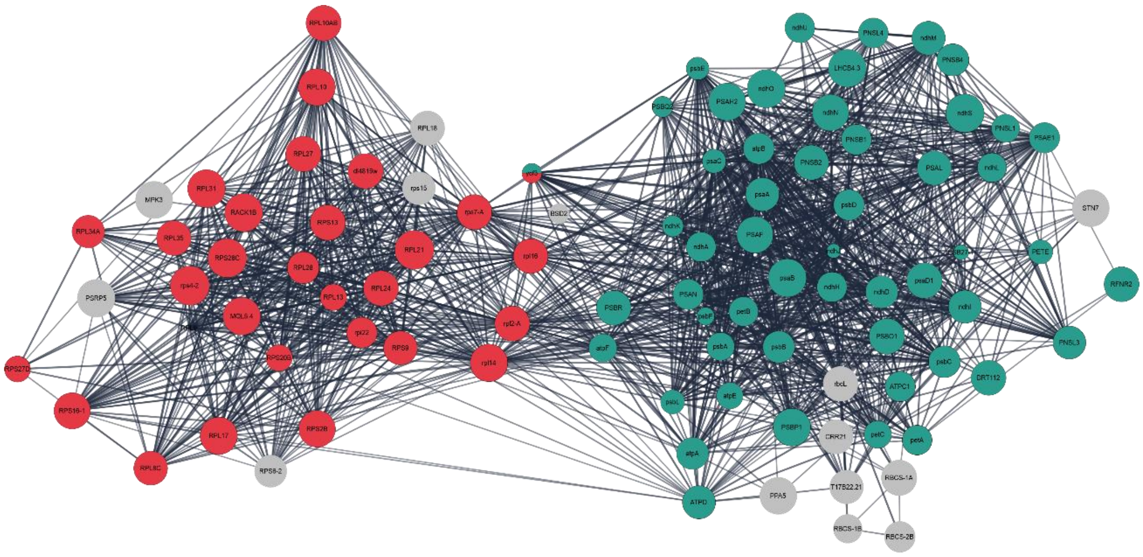

**Supplemental Figure 4: Interaction network of downregulated proteins in LIN-treated seedlings.** Proteins with lowered abundances in response to 15 µM LIN treatment were mapped using STRING interaction data. Downregulated proteins ($log_2$ fold change ≤ –1; node size reflects magnitude) formed a highly interconnected network with two major clusters: ribosomal proteins (red nodes) and proteins associated with photosynthesis (blue-green nodes).
